# Foot placement coordination is impaired in people with Parkinson’s Disease

**DOI:** 10.1101/2025.06.11.659042

**Authors:** Charlotte Lang, Jeffrey M Hausdorff, Sjoerd M Bruijn, Matthew A Brodie, Yoshiro Okubo, Walter Maetzler, Moira van Leeuwen, Navrag B Singh, Jaap H van Dieen, Deepak K Ravi

## Abstract

**Background:** Gait instability is a common and disabling symptom of Parkinson’s disease (PD), contributing to frequent falls and reduced quality of life. While clinical balance tests and spatiotemporal gait measures can predict fall risk, they do not fully explain the underlying control mechanisms. In healthy individuals, foot placement is actively adjusted based on an internal estimate of the center of mass state to maintain gait stability, known as foot placement control. This estimation relies on the integration of multisensory information, which has been shown to be impaired in PD, potentially disrupting the control of gait stability through foot placement.

**Objective:** To investigate whether foot placement coordination during overground walking is impaired in people with PD.

**Methods:** Fifty people with PD and 51 healthy older adults walked overground for ten minutes at self-selected, comfortable walking speed. Foot placement errors were quantified as the deviation between the actual foot placement and the predicted placement derived from the center of mass kinematic state during the preceding swing phase.

**Results:** Foot placement errors were significantly higher in people with PD than in healthy older adults in both mediolateral (p < .05) and anteroposterior directions (p < .0001), at both mid-swing and terminal swing. Relative explained variance in mediolateral direction was significantly higher in people with PD compared to healthy older adults (p < .005).

**Conclusion:** We provide first evidence of impaired coordination between the center of mass and foot placement in PD. Future work should investigate a causal relationship between impaired foot placement control, sensorimotor integration and gait instability.

## Introduction

Gait instability is a common and disabling symptom of Parkinson’s disease (PD). People with PD often walk slowly with short, shuffling and unstable steps, which increases their risk of falling. Up to 50% of individuals with PD experience recurrent falls, contributing significantly to reduced quality of life [1-3]. Several clinical tests have been validated and are widely used to assess balance performance in PD [4]. These include the BESTest and the miniBESTest [5, 6], both of which are known for their strong predictive value in distinguishing fallers from non-fallers. Other commonly used assessments include the Functional Gait Assessment [7], which assesses dynamic balance during walking, and the Berg-Balance-Scale [8, 9], which focuses exclusively on static balance.

Beyond clinical assessments, laboratory-based measures, such as walking speed, step length, and stride time variability are frequently used and have been shown to differ significantly in people with PD relative to healthy adults [10, 11]. These gait characteristics are significantly associated with fall risk in PD [12, 13]. For instance, slower gait speed and higher variability in step timing have been identified as strong predictors of future falls in individuals with PD [12]. However, while these metrics reflect aspects of gait instability, they do not fully explain the underlying control mechanisms, which therefore remain poorly understood in PD. Identifying which stability control mechanisms are impaired in PD may help guide the development and improvement of interventions targeting gait instability.

Healthy individuals stabilize steady-state walking, meaning in the absence of large external disturbances or changes in gait speed, primarily using a strategy known as foot placement control. During each step, foot placement is adjusted based on the estimated state of the Center of Mass (CoM), specifically its position and velocity, during the preceding swing phase [14]. This allows for deviations in CoM movement from a stable trajectory to be accommodated by actively stepping in the direction of the disturbance. Here, the degree of foot placement control can be quantified by how well foot placement can be predicted from the CoM kinematic state [15]. In healthy young adults, this relationship is strong, with CoM dynamics explaining over 80% of the variance in mediolateral foot placement, highlighting its relevance for maintaining balance during walking [15]. However, in older adults and individuals with stroke, the degree of foot placement control has been shown to be diminished [16, 17], highlighting its susceptibility to age- and disease-related changes in sensorimotor function.

Accurate estimation of the CoM state and appropriate foot placement adjustments rely on the integration of sensory information from the vestibular, visual and proprioceptive systems [14, 18, 19]. In people with PD, impairments in sensory processing and sensorimotor integration have been previously reported [20-23], which may compromise CoM state estimation and reduce the precision of foot placement. This, in turn, could lead to greater gait instability and an increased risk of falling. Therefore, this study aims to investigate whether foot placement control is impaired during overground walking in people with PD.

## Methods

### Dataset

This study re-analyses an existing dataset from the Laboratory for Movement Biomechanics (ETH Zurich), previously collected and presented by Mei et al. [24]. The dataset includes 50 individuals with PD (41 male, mean age: 60 (SD 11) years, mean PD duration: 9 (5) years, mean MDS-UPDRS III in ON medication: 19 (7)) and 51 healthy controls without any known diseases or conditions (22 male, mean age: 67 (11) years) (Table 1). One of the main inclusion criteria required participants to be able to walk independently and continuously for 10 minutes. All assessments of the PD participants were performed during ON-medication state. Participants walked barefoot continuously for 10 minutes at a self-selected, comfortable speed without assistance, following a path that resembled an elongated figure-8: two 10-meter segments connected by large-radius turns around two signposts. Only the straight walking segments were recorded and included in the analysis.

**Table 1:**
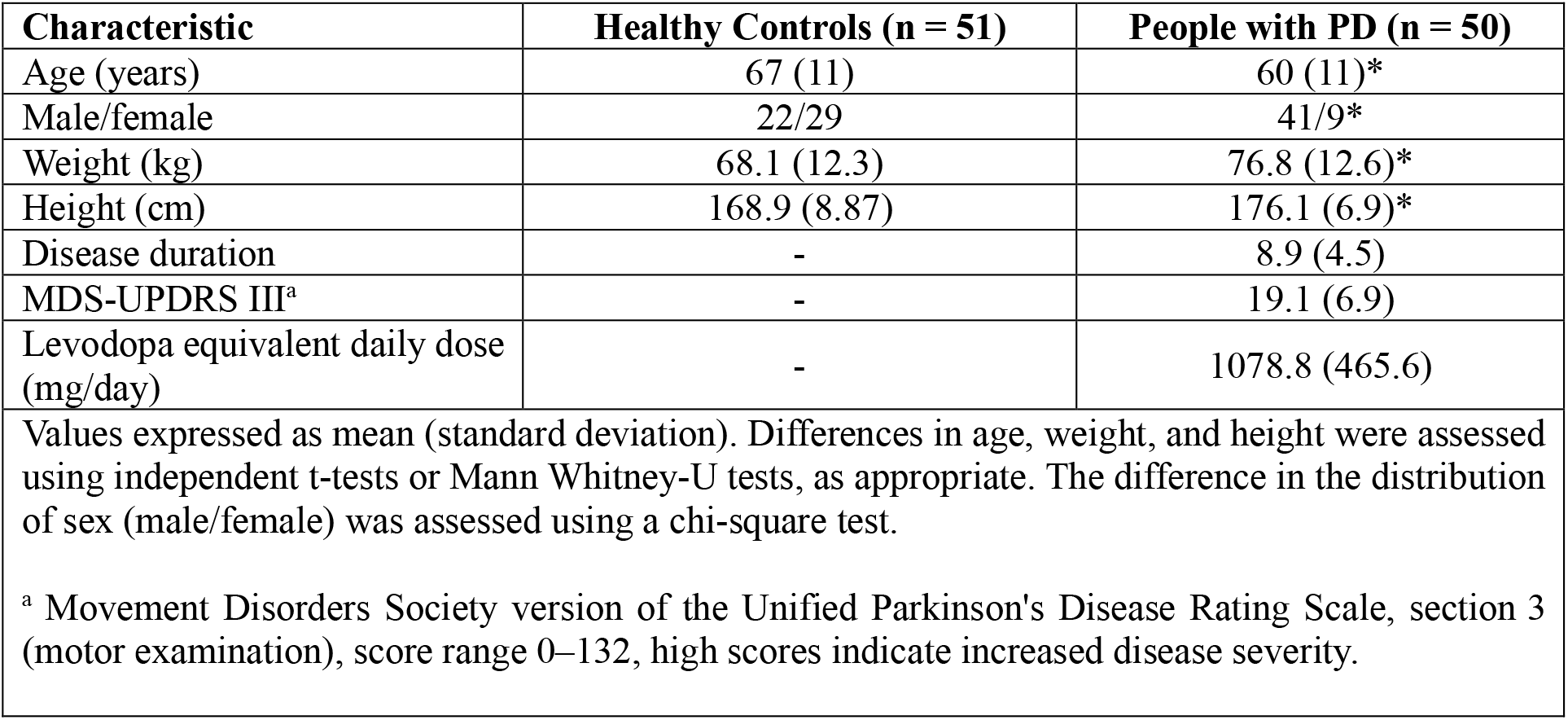
Demographics and clinical characteristics. Significant differences between groups are indicated by * (p < .05)

### Data collection, processing and analysis

Full body kinematics data were collected at a sample rate of 100 Hz using an optical motion capture system (61 markers, Vicon Nexus, version 2.3/2.8.2, Oxford Metrics, United Kingdom). Gait events (heel-strike and toe-off) were detected using a custom algorithm based on velocity of the foot markers [25]. The sacrum marker was used to estimate mediolateral and anteroposterior CoM position. The heel marker was used to extract foot trajectories. Swing phases were identified based on the detected toe-off and heel-strike events and subsequently time-normalized to 0-100% using spline interpolation. This process was performed for both mediolateral and anteroposterior movement direction. The last step of each straight path segment was removed and an array containing all valid steps was then generated for the analysis. For each straight path segment, between 5 and 8 steps remained for the analysis.

### Foot placement model

Foot placement control is reflected in step width (i.e. the mediolateral distance between heel markers, where the position of each foot was determined at their respective midstance, ensuring both feet are flat on the ground) and step length (i.e. the anteroposterior distance at heelstrike of the leading foot) which are predicted from the CoM kinematic state during swing phase of walking using linear models. Each time-normalized swing phase results in 51 equally spaced samples, representing a specific time point within the swing phase. Mediolateral and anteroposterior CoM position and velocity at each of these samples were used as predictors of foot placement. CoM displacement was calculated with respect to the stance foot, and CoM velocity as its derivative. All variables were demeaned before fitting the models for mediolateral (step width, SW) and anteroposterior direction (step length, SL):

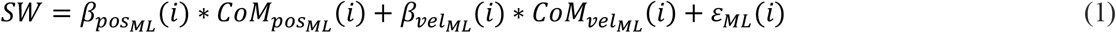

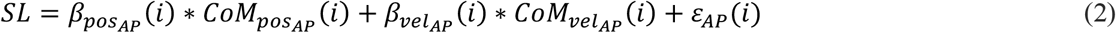

The regression coefficients (β) represent the “strength” of the foot placement response to CoM state deviations from the mean. The Root Mean Square of the residuals (ε) is referred to as foot placement error (FPE), indicating the precision of foot placement control. Each sample (i) of the swing phase represents a different normalized time point of the swing phase. Model outcomes were evaluated at two points during the swing phase: mid-swing (i = 25) and terminal swing (i = 51). Mid-swing is a point where the system can still adjust the leg’s trajectory based on sensory feedback, so it reflects feedback control. Terminal swing occurs just before the foot hits the ground (heel strike), so it reflects the final outcome of the control process, how accurately the foot is placed to maintain stability. Relative explained variance (R^2^) was determined as the variance in actual foot placement accounted for by the predicted foot placement, representing the degree of foot placement control.

### Statistics

For the comparison of demographics (age, height, weight), walking speed and number of steps between people with PD and healthy older adults, either t-tests or Mann Whitney-U tests were performed, depending on the normality of the data distribution. Difference in number of male/female participants was assessed using a chi-square test. As the primary analysis, ANCOVAs were performed to test whether foot placement error and relative explained variance (dependent variables) were affected by the independent variables Group (PD vs. Control) and Timepoint (mid-swing vs. terminal swing), while adjusting for step length, step width and age as covariates. Gait speed strongly covaries with step length and was therefore not included as an additional covariate [26]. Separate analyses were conducted for mediolateral and anteroposterior directions. Data normality was assessed using the Shapiro-Wilk test. In case of a skewed distribution, data of foot placement error were log transformed. Relative explained variance was transformed using modified a Fisher transformation. Post hoc group comparisons at each timepoint were performed using Tukey’s correction for multiple comparisons. As a secondary analysis, foot placement variability, i.e. how much the position of the foot varies relative to the stance foot, and CoM variability was compared between groups using Mann Whitney-U tests. *P* values < .05 were considered statistically significant for all analyses.

All analyses were conducted using Matlab (version R2023b, The MathWorks Inc., Natick) and R (v4.3.1, The R Foundation for Statistical Computing, Austria)

## Results

The average gait speed was 1.20 (0.19) m/s in people with PD and 1.21 (0.14) m/s in healthy controls (p = .75). The average number of steps included in the analysis was 413 ± 123 for the PD group and 354 ± 121 for the control group (p < .005). There were significant differences (p < .05) in age, sex distribution, height, and weight between people with PD and healthy controls. Given the difference in sex distribution, a sub-analysis was conducted to examine potential sex-related effects on foot placement outcomes (Supplementary material). However, no significant differences between male and female participants were found and therefore this was not further considered in the analysis.

### Foot placement error

In the mediolateral direction, a significant main effect of both Group (PD vs. control; p < .05) and Timepoint (mid-swing vs. terminal swing; p < .0001) was found, while there was no significant Group × Timepoint interaction (p = .70). This indicates that, even after adjusting for step length, step width, and age, people with PD had consistently higher foot placement errors across timepoints. Post hoc comparison confirmed that foot placement error was significantly higher in the PD group compared to controls at both mid-swing (p < .05) and terminal swing (p = .005) (Figure 1, left). Among the covariates, longer step length was associated with higher mediolateral foot placement error (p < .005).

**Figure 1:**
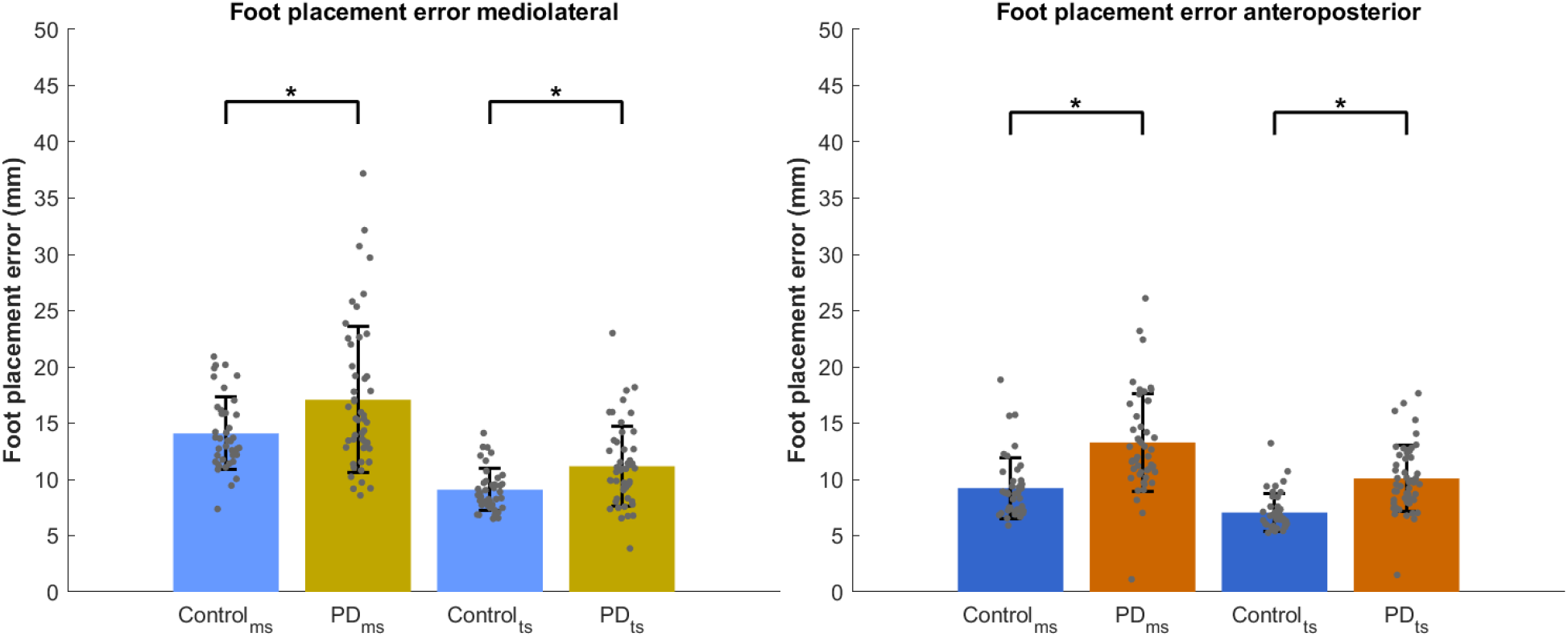
Foot placement errors (mm) for people with PD and healthy controls during mid-swing (ms) and terminal swing (ts) in both mediolateral (left) and anteroposterior (right) direction. Error bars represent standard deviation.

In the anteroposterior direction, there were also significant main effects of Group (p < .0001) and Timepoint (p < .0001), but no significant interaction (p = .94). Post hoc comparison showed that the PD group had significantly higher foot placement errors than controls at both mid-swing and terminal swing (p < .0001) (Figure 1, right). Of the covariates, larger step widths were significantly associated with increased anteroposterior foot placement error (p < .001).

### Relative explained variance (R2)

In the mediolateral direction, a significant main effect of Group (p < .005) and Timepoint (p < .0001) was observed after adjusting for step width, step length and age. The Group × Timepoint interaction was not significant (p = .57), indicating that the effect of group was consistent across both timepoints. CoM state at mid-swing explained 82% of the variance in subsequent foot placement in people with PD compared to 78% in the control group (p < .005) (Figure 2, left). The same held for terminal swing, where the CoM state accounted for significantly more foot placement variance in PD (93% vs. 91% in controls, p < .05). The outcome was significantly influenced by step width (p < .001), with wider steps being associated with higher R^2^.

**Figure 2:**
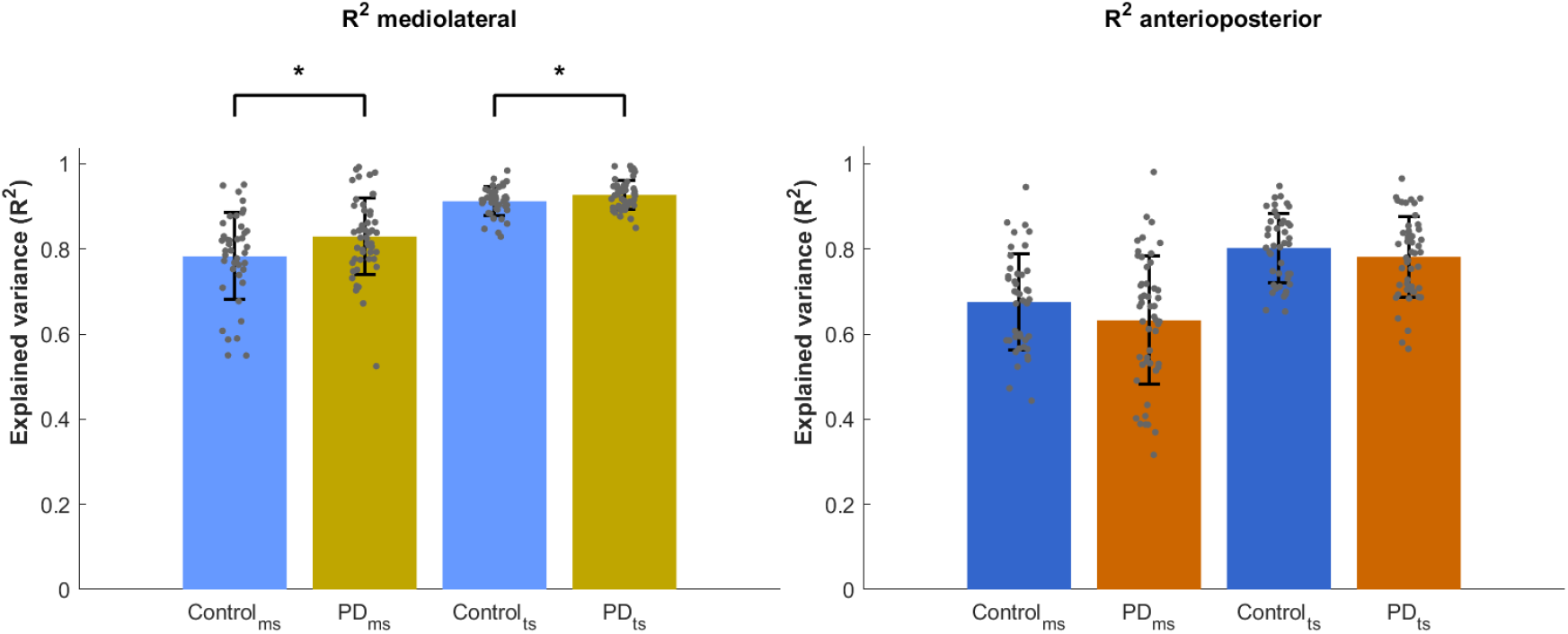
Relative explained variance (R^2^) for people with PD and healthy controls at mid-swing (ms) and terminal swing (ts) for both mediolateral (left) and anteroposterior (right) direction. Error bars represent standard deviation.

In the anteroposterior direction, the CoM state at mid-swing explained 63% of the variance in foot placement in people with PD and 67% in controls, increasing to 78% and 80%, respectively, at terminal swing (Figure 2, right). No main effect of group or Group × Timepoint interaction was found, but there was a significant main effect of Timepoint (p < .001), indicating that the increase in R^2^ from mid-swing to terminal swing occurred similarly in both groups.

### Secondary outcomes

To test whether changes in R^2^ were (partially) due to increases in variability [27], differences in foot placement and CoM variability were tested, showing a significantly higher variability of step width/step length and CoM position in PD compared to controls for both ML and AP (p < .001) (Table 2).

**Table 2:** Center of mass and foot placement variability (mm) for both mediolateral and anteroposterior direction. Significant differences between groups are indicated by * (p < .05)

|  | Healthy Controls | People with PD |
| --- | --- | --- |
| CoM variability ML (mm) | 15.28 (3.69) | 21.72 (10.55)* |
| CoM variability AP (mm) | 13.28 (3.28) | 15.55 (3.56)* |
| SW variability (mm) | 42.78 (19.29) | 49.15 (20.68)* |
| SL variability (mm) | 20.76 (7.31) | 29.61 (11.45)* |

## Discussion

This study investigated foot placement control during overground walking in people with PD compared to healthy older adults. Our results revealed that people with PD exhibit significantly higher foot placement errors in both the mediolateral and anteroposterior directions, at both mid-swing and terminal swing phases. Relative explained variance in mediolateral direction is increased compared to healthy older adults. Together, our study documents novel aspects of impaired foot placement control (feedback control) in people with PD.

People with PD had higher foot placement errors than healthy older adults. This may result from either an impairment in the integration of sensory modalities or impaired neural control of gait. An impaired integration of sensory modalities could potentially lead to an imprecise estimation of the CoM state [22]. As a result, people with PD may place their feet less precisely, limiting their ability to adequatly compensate deviations in CoM movement through appropriate foot placement. Such impaired adaptatation could contribute to reduced stability and an increased risk of falls. An impaired neural control of gait in PD can result from changes in the beta band, which play a crucial role for gait control [28, 29]. Beta oscillations in the sensorimotor cortex have long been recognized as key signatures of movement, closely linked to sensorimotor integration control [30]. For example, in healthy adults, beta activity is involved in stabilization of gait [31]. In PD the function of beta activity is altered and influenced by levodopa, leading to an impaired of neural gait control leading to less sabilized gait [32]. In addition to sensory and neural deficits, biomechanical factors may also contribute to impaired foot placement. Previous studies have shown that people with PD exhibit altered foot strike angles and reduced toe clearance at the end of the swing phase compared to healthy older adults [33]. These changes may reflect impaired motor control of the swing leg, possibly influenced by the same sensorimotor deficits, and could constrain the ability to execute precise foot placement. However, such associations remain to be investigated.

The findings of this study demonstrate a difference in foot placement error of approximately 3 to 4 mm between people with PD and healthy older adults. While the absolute difference may seem modest, it represents a substantial relative change given that typical foot placement precision ranges from 8 to 15 mm, highlighting the potential clinical relevance of even small disruptions in feedback control. Furthermore, Magnani et al. investigated how vestibular stimulation affects gait stability and found that even small differences in foot placement error between similar conditions were associated with reduced stability [27]. Despite the absence of a direct causal relationship being investigated, this emphasises the critical role of precise foot placement in maintaining stable gait. Therefore, the differences in foot placement precision evident in this study may bear significant clinical implications, as even modest reductions in precision might have the potential to compromise gait stability and elevate the risk of falling.

Regarding the relative explained variance in mediolateral direction, people with PD exhibited a higher degree of foot placement control (indiciated by a higher R^2^), suggesting that mediolateral foot placement in PD may be more strongly governed by CoM state than in healthy adults. This would indicate a more tightly regulated foot placement based on CoM state in the mediolateral direction among people with PD. However, R^2^ is influenced by the signal-to-noise ratio and the increased variability in foot placement and CoM position, which we observed in our data and is common in PD, can increase errors and inflate R^2^ values. Therefore, the elevated explained variance in the PD group may not necessarily indicate enhanced control, but could instead reflect greater movement variability. On the other hand, there may also exist an optimal level of control, beyond which movement may be more likely to become rigid, with tighter control potentially leading to an inability to adapt to disturbances [34]. During gait, the moving base of support is narrowest in the ML direction [35]. In the single-limb stance phase, the center of mass moves closest to the lateral edge of the base of support and therefore regulation in the mediolateral direction may be challenging [14]. People with PD may compensate for this stability challenge by tighter control of ML foot placements, indicated by an increased explained variance, and decreased degrees of movement freedom. However, it remains to be investigated whether this increase is due to methodological issues arising from higher variability or indeed a higher level of control, whether beneficial or detrimental, in people with PD. No difference was found between people with PD and healthy older adults in anteroposterior direction.

In the dataset used, falls were not systematically recorded, and therefore, the relationship between foot placement and falls could not be thoroughly investigated. However, self-reported fall histories obtained from their medical history were available for some participants with PD. We examined foot placement errors in three individual cases who reported frequent falls: (a) one fall per week, (b) 3–4 falls in the previous month, and (c) a total of 10 falls. All three cases showed increased foot placement errors at terminal swing, with the largest errors observed in the participant reporting one fall per week (ML: 23.0 mm, AP: 16.1 mm). These findings suggest a potential link between impaired foot placement control and fall risk and highlight the need for future studies to record falls systematically.

Although we ensured that the overground walking steps were straight ahead (the curved parts of the figure 8 shapes were excluded), we cannot ascertain that participants did not already adjust their steps in anticipation of the turn. Curved and straight line walking differ in several aspects, such as asymmetries between the inside and outside leg, slowing down or mediolateral displacement of the CoM during turning [36]. To minimize such potential effects the last step of each straight walking path was excluded. However, given the short walking distance it was not possible to remove more than one step at the end of each path. Therefore, some included steps may not reflect steady-state walking, potentially affecting the foot placement outcomes.

In conclusion, our results provide evidence that feedback control is impaired in people with PD, as indicated by a less precise coordination between foot placement and CoM kinematics, which appears to be consistent across both anteroposterior and mediolateral movement directions. Given the association of foot placement control with gait stability [14], the present results may be an early step toward a mechanistic understanding of gait instability in PD, which is important for the development of effective approaches in gait rehabilitations. Designing interventions that target foot placement control and increase the precision of foot placement might improve gait stability in people with PD. Further research is needed to identify how impaired sensorimotor integration in PD affects the successful execution of this foot placement strategy.

## Aknowledgements

The authors would like to thank all project members involved in the original data collection, including Prof. Dr. William Taylor, Michelle Gwerder, Elena Bernasconi, Giancarlo Crameri, Ashwak Hussein, Angela Frautschi and Niklas König Ignasiak from ETH Zurich as well as Dr. Christian R. Baumann, Mechtild Uhl and Lennart Stieglitz from University Hospital Zurich. The authors further would like to thank Dr. Julius Welzel for his support to this work.

## References

1. Ebersbach, G., C. Moreau, F. Gandor, L. Defebvre, and D. Devos, Clinical syndromes: Parkinsonian gait. Mov Disord, 2013. 28(11): p. 1552–9.

2. Fasano, A., C.G. Canning, J.M. Hausdorff, S. Lord, and L. Rochester, Falls in Parkinson’s Disease: A Complex and Evolving Picture. Movement Disorders, 2017. 32(11): p. 1524–1536.

3. Pelicioni, P.H.S., J.C. Menant, M.D. Latt, and S.R. Lord, Falls in Parkinson’s Disease Subtypes: Risk Factors, Locations and Circumstances. International Journal of Environmental Research and Public Health, 2019. 16(12).

4. Bloem, B.R., J. Marinus, Q. Almeida, L. Dibble, A. Nieuwboer, B. Post, et al., Measurement instruments to assess posture, gait, and balance in Parkinson’s disease: Critique and recommendations. Movement Disorders, 2016. 31(9): p. 1342–1355.

5. Horak, F.B., D.M. Wrisley, and J. Frank, The Balance Evaluation Systems Test (BESTest) to Differentiate Balance Deficits. Physical Therapy, 2009. 89(5): p. 484–498.

6. King, L.A., K.C. Priest, A. Salarian, D. Pierce, and F.B. Horak, Comparing the Mini-BESTest with the Berg Balance Scale to Evaluate Balance Disorders in Parkinson’s Disease. Parkinsons Disease, 2012. 2012.

7. Leddy, A.L., B.E. Crowner, and G.M. Earhart, Functional Gait Assessment and Balance Evaluation System Test: Reliability, Validity, Sensitivity, and Specificity for Identifying Individuals With Parkinson Disease Who Fall. Physical Therapy, 2011. 91(1): p. 102–113.

8. Qutubuddin, A.A., P.O. Pegg, D.X. Cifu, R. Brown, S. McNamee, and W. Carne, Validating the Berg Balance Scale for patients with Parkinson’s disease: A key to rehabilitation evaluation. Archives of Physical Medicine and Rehabilitation, 2005. 86(4): p. 789–792.

9. Steffen, T. and M. Seney, Test-retest reliability and minimal detectable change on balance and ambulation tests, the 36-Item Short-Form Health Survey, and the unified Parkinson disease rating scale in people with parkinsonism. Physical Therapy, 2008. 88(6): p. 733–746.

10. Zanardi, A.P.J., E.S. da Silva, R.R. Costa, E. Passos-Monteiro, I.O. dos Santos, L.F.M. Kruel, et al., Gait parameters of Parkinson’s disease compared with healthy controls: a systematic review and meta-analysis. Scientific Reports, 2021. 11(1).

11. Mirelman, A., P. Bonato, R. Camicioli, T.D. Ellis, N. Giladi, J.L. Hamilton, et al., Gait impairments in Parkinson’s disease. Lancet Neurology, 2019. 18(7): p. 697–708.

12. Lord, S., B. Galna, A.J. Yarnall, S. Coleman, D. Burn, and L. Rochester, Predicting First Fall in Newly Diagnosed Parkinson’s Disease: Insights From a Fall-Naive Cohort. Movement Disorders, 2016. 31(12): p. 1829–1836.

13. Creaby, M.W. and M.H. Cole, Gait characteristics and falls in Parkinson’s disease: A systematic review and meta-analysis. Parkinsonism & Related Disorders, 2018. 57: p. 1–8.

14. Bruijn, S.M. and J.H. van Dieën, Control of human gait stability through foot placement. Journal of the Royal Society Interface, 2018. 15(143).

15. Wang, Y. and M. Srinivasan, Stepping in the direction of the fall: the next foot placement can be predicted from current upper body state in steady-state walking. Biology Letters, 2014. 10(9).

16. Dean, J.C. and S.A. Kautz, Foot placement control and gait instability among people with stroke. Journal of Rehabilitation Research and Development, 2015. 52(5): p. 577–590.

17. Arvin, M., M.J.M. Hoozemans, M. Pijnappels, J. Duysens, S.M. Verschueren, and J.H. van Dieen, Where to Step? Contributions of Stance Leg Muscle Spindle Afference to Planning of Mediolateral Foot Placement for Balance Control in Young and Old Adults. Front Physiol, 2018. 9: p. 1134.

18. O’Connor, S.M. and A.D. Kuo, Direction-Dependent Control of Balance During Walking and Standing. Journal of Neurophysiology, 2009. 102(3): p. 1411–1419.

19. van Schooten, K.S., L.H. Sloot, S.M. Bruijn, H. Kingma, O.G. Meijer, M. Pijnappels, et al., Sensitivity of trunk variability and stability measures to balance impairments induced by galvanic vestibular stimulation during gait. Gait & Posture, 2011. 33(4): p. 656–660.

20. Hwang, S., P. Agada, S. Grill, T. Kiemel, and J.J. Jeka, A central processing sensory deficit with Parkinson’s disease. Experimental Brain Research, 2016. 234(8): p. 2369–2379.

21. Smith, P.F., Vestibular Functions and Parkinson’s Disease. Frontiers in Neurology, 2018. 9.

22. Kearney, J. and J.S. Brittain, Sensory Attenuation in Sport and Rehabilitation: Perspective from Research in Parkinson’s Disease. Brain Sciences, 2021. 11(5).

23. Feller, K.J., R.J. Peterka, and F.B. Horak, Sensory Re-weighting for Postural Control in Parkinson’s Disease. Frontiers in Human Neuroscience, 2019. 13.

24. Mei, Z.H., A. S.; Baumann, C; Uhl, M.; Singh, N.; Taylor, W.R.; Stieglitz, L.; Ravi, D. K., The Role of Electrode Placement in STN-DBS for Improving Gait in Parkinson’s Disease. medRxiv, 2023.

25. O’Connor, C.M., S.K. Thorpe, M.J. O’Malley, and C.L. Vaughan, Automatic detection of gait events using kinematic data. Gait & Posture, 2007. 25(3): p. 469–474.

26. Collins, S.H. and A.D. Kuo, Two Independent Contributions to Step Variability during Over-Ground Human Walking. Plos One, 2013. 8(8).

27. Magnani, R.M., J.H. van Dieën, and S.M. Bruijn, Effects of vestibular stimulation on gait stability when walking at different step widths. Experimental Brain Research, 2023. 241(1): p. 49–58.

28. Roeder, L., T.W. Boonstra, and G.K. Kerr, Corticomuscular control of walking in older people and people with Parkinson’s disease. Scientific Reports, 2020. 10(1).

29. Jacobsen, N.S.J., S. Blum, J.E.M. Scanlon, K. Witt, and S. Debener, Mobile electroencephalography captures differences of walking over even and uneven terrain but not of single and dual-task gait. Front Sports Act Living, 2022. 4: p. 945341.

30. Tan, H.L., C. Wade, and P. Brown, Post-Movement Beta Activity in Sensorimotor Cortex Indexes Confidence in the Estimations from Internal Models. Journal of Neuroscience, 2016. 36(5): p. 1516–1528.

31. Bruijn, S.M., J.H. Van Dieën, and A. Daffertshofer, Beta activity in the premotor cortex is increased during stabilized as compared to normal walking. Frontiers in Human Neuroscience, 2015. 9.

32. Tinkhauser, G., A. Pogosyan, H.L. Tan, D.M. Herz, A.A. Kühn, and P. Brown, Beta burst dynamics in Parkinson’s disease OFF and ON dopaminergic medication. Brain, 2017. 140: p. 2968–2981.

33. Ginis, P., R. Pirani, S. Basaia, A. Ferrari, L. Chiari, E. Heremans, et al., Focusing on heel strike improves toe clearance in people with Parkinson’s disease: an observational pilot study. Physiotherapy, 2017. 103(4): p. 485–490.

34. Ravi, D.K., M. Gwerder, N.K. Ignasiak, C.R. Baumann, M. Uhl, J.H. van Dieën, et al., Revealing the optimal thresholds for movement performance: A systematic review and meta-analysis to benchmark pathological walking behaviour. Neuroscience and Biobehavioral Reviews, 2020. 108: p. 24–33.

35. Curtze, C., T.J.W. Buurke, and C. McCrum, Notes on the margin of stability. Journal of Biomechanics, 2024. 166.

36. Guglielmetti, S., A. Nardone, A.M. De Nunzio, M. Godi, and M. Schieppati, Walking Along Circular Trajectories in Parkinson’s Disease. Movement Disorders, 2009. 24(4): p. 598–604.

