## Supplementary Material for "Foot placement coordination is impaired in people with Parkinson’s Disease"

### Foot placement error mediolateral

| **Effect** | | **p Mid-swing** | | **p Terminal swing** |
| --- | --- | --- | --- | --- |
| Group |  | **0.0367** |  | **0.0067** |
| Gender |  | 0.9745 |  | 0.9909 |
| SW |  | 0.7332 |  | 0.4922 |
| SL |  | 0.1538 |  | 0.0717 |
| Age |  | 0.3735 |  | 0.3463 |
| Group * Gender |  | 0.1259 |  | 0.0914 |

### Foot placement error anteroposterior

| **Effect** | | **p Mid-swing** | | **p Terminal swing** |
| --- | --- | --- | --- | --- |
| Group |  | **1.9e-05** |  | **7.52e-06** |
| Gender |  | 0.4666 |  | 0.5638 |
| SW |  | 0.0193 |  | 0.0235 |
| SL |  | 0.8511 |  | 0.4603 |
| Age |  | 0.3084 |  | 0.9836 |
| Group * Gender |  | 0.0613 |  | 0.0741 |

### Explained variance mediolateral

| **Effect** | | **p Mid-swing** | | **p Terminal swing** |
| --- | --- | --- | --- | --- |
| Group |  | **0.0136** |  | **0.0162** |
| Gender |  | 0.4984 |  | 0.5120 |
| SW |  | **0.0002** |  | **2.54e-05** |
| SL |  | 0.8040 |  | 0.5238 |
| Age |  | 0.3900 |  | 0.2670 |
| Group * Gender |  | 0.3426 |  | 0.3039 |

### Explained variance anteroposterior

| **Effect** | | **p Mid-swing** | | **p Terminal swing** |
| --- | --- | --- | --- | --- |
| Group |  | 0.1045 |  | 0.4807 |
| Gender |  | 0.9719 |  | 0.8610 |
| SW |  | 0.9756 |  | 0.4502 |
| SL |  | 0.4181 |  | 0.7007 |
| Age |  | 0.6287 |  | **0.0073** |
| Group * Gender |  | 0.4074 |  | 0.6308 |
